# Self-repair protects microtubules from their destruction by molecular motors

**DOI:** 10.1101/499020

**Authors:** Sarah Triclin, Daisuke Inoue, Jérémie Gaillard, Zaw Min Htet, Morgan E. DeSantis, Didier Portran, Emmanuel Derivery, Charlotte Aumeier, Laura Schaedel, Karin John, Christophe Leterrier, Samara L. Reck-Peterson, Laurent Blanchoin, Manuel Théry

**Affiliations:** Univ. Grenoble-Alpes, CEA, CNRS, INRA, Biosciences & Biotechnology Institute of Grenoble, Laboratoire de Phyiologie Cellulaire & Végétale, CytoMorpho Lab, 38054 Grenoble, France; Deptartment of Cellular and Molecular Medicine, and Cell and Developmental Biology Section, Division of Biological Sciences, University of California San Diego, 9500 Gilman Drive, La Jolla, CA 92093, USA; Howard Hughes Medical Institute; CRBM - CNRS UMR 5237, Route de Mende, 34293 Montpellier Cedex 05, FRANCE; MRC Laboratory of Molecular Biology, Francis Crick Avenu, Cambridge Biomedical Campus, Cambridge CB2 0QH, UK; University of Geneva Department of Biochemistry, Science II 30, Quai Ernest-Ansermet 1205 Genève, Switzerland; Univ. Grenoble-Alpes, CNRS, Laboratoire Interdisciplinaire de Physique, 38000 Grenoble, France; Aix Marseille Université, CNRS, INP UMR7051, Marseille, France; Univ. Paris Diderot, INSERM, CEA, Hôpital Saint Louis, Institut Universitaire d’Hematologie, UMRS1160, CytoMorpho Lab, 75010 Paris, France

**Author notes:** these authors contributed equally to this work (Sarah Triclin, Daisuke Inoue).

## Abstract

Microtubules are dynamic polymers that are used for intracellular transport and chromosome segregation during cell division. Their instability stems from the low energy of tubulin dimer interactions, which sets the growing polymer close to its disassembly conditions^1^. Microtubules function in coordination with kinesin and dynein molecular motors^2–4^, which use ATP hydrolysis to produce mechanical work and move on microtubules^5^. This raises the possibility that the forces produced by walking motors can break dimer interactions and trigger microtubule disassembly. We tested this hypothesis by studying the interplay between microtubules and moving molecular motors in vitro. Our results show that the mechanical work of molecular motors can remove tubulin dimers from the lattice and rapidly destroy microtubules. This effect was not observed when free tubulin dimers were present in the assay. Using fluorescently labelled tubulin dimers we found that dimer removal by motors was compensated for by the insertion of free tubulin dimers into the microtubule lattice. This self-repair mechanism allows microtubules to survive the damage induced by molecular motors as they move along their tracks. Our study reveals the existence of coupling between the motion of kinesin and dynein motors and the renewal of the microtubule lattice.

---

Molecular motors can alter microtubule growth and shrinkage^6^. Kinesins impact microtubule growth by regulating the dynamics of dimer addition or the recruitment of macro-molecular complexes at microtubule ends^6,7^. Dyneins, the other family of microtubule-based motors, can capture microtubule plus ends and induce slow microtubule depolymerisation^8^. The kinesin-13 and kinesin-8 families actively depolymerise microtubules by removing dimers at microtubule ends^9,10^. These motors travel along the shaft and modulate microtubule dynamics as they reach microtubule ends, but have not been shown to have any impact on the microtubule shaft. However, thermal forces are sufficient to promote the removal of tubulin dimers from the lattice^11^ and motors locked into a non-moving state are capable of expanding the microtubule lattice^12,13^, suggesting that the mechanical work produced by moving motors could impact the stability and dynamics of microtubule shaft.

An analysis of the energy performance of walking motors suggests that they could indeed break the bonds between tubulin dimers. Molecular motors use ATP hydrolysis to drive conformational changes, which powers their movement along the microtubule shaft^14,15^. For example, kinesins hydrolyze one ATP (about 22 kT) per 8 nm step^5^ to produce a maximal stall force of about 7 pN^16^. Thus, in the most efficient scenario the mechanical work produced by kinesin is close to 56 pN.nm (14 kT), suggesting that 8 kT are dissipated with each motor step. Moreover, ATP hydrolysis does not power the entire step^17^, but rather only the neck-linker docking step resulting in a smaller 4 nm forward displacement of the rear head that corresponds to 24 pN.nm (6 kT)^14,18^. This calculation suggests that 16 kT are dissipated during each kinesin step^19,20^. The estimated lateral and longitudinal bond energies linking tubulin dimers within the microtubule lattice are ~ 5kT and 23 kT, respectively^21^. Thus walking motors dissipate sufficient energy to disrupt the microtubule structure.

Most in vitro studies of the mechanical forces produced by molecular motors have been performed on stabilised microtubules in order to block microtubule depolymerisation^22^. Indeed, under physiological conditions GTP-tubulin is rapidly hydrolysed to GDP-tubulin following tubulin dimer incorporation at the growing microtubule end. Because the GDP-tubulin lattice is less stable than that with GTP-tubulin rapid disassembly can occur starting from microtubule ends in assays on non-stabilized microtubules. However, this unstable state corresponds to the actual conformation of the shaft of microtubules in vivo. Here we set out to study the impact of mechanical forces produced by kinesin and dynein motors on microtubules under conditions where the lattice would be made up of GDP-tubulin. We started by using microtubule "gliding assays”, where motors are attached to the glass surface of a small flow cell. Upon the addition of microtubules and ATP the forces produced by active motors results in microtubule gliding across the motor-coated surface.

We first performed gliding assays with the plus-end-directed human kinesin-1 on microtubules stabilized either with taxol (Figure 1A-i) or by assembling microtubules with GMPCPP-bound tubulin, which traps tubulin in a state that mimics stable GTP-bound tubulin (Figure A-ii). The total amount of microtubules did not change during the time-course of the assay in either condition (Figure 1B, 1C, movie S1). Next, to study the response of GDP-lattice microtubules, we capped microtubules on both ends with a short microtubule section consisting of GMPCPP-tubulin, which protects microtubules from depolymerising from their ends (Figure 1A-iii, see Methods). These capped-GDP microtubules started depolymerizing within minutes of initiating the gliding assay (Figure 1B, movie S2). Fifteen minutes later, no microtubules were detectable on the slide (Figure 1C).

**Figure 1.**
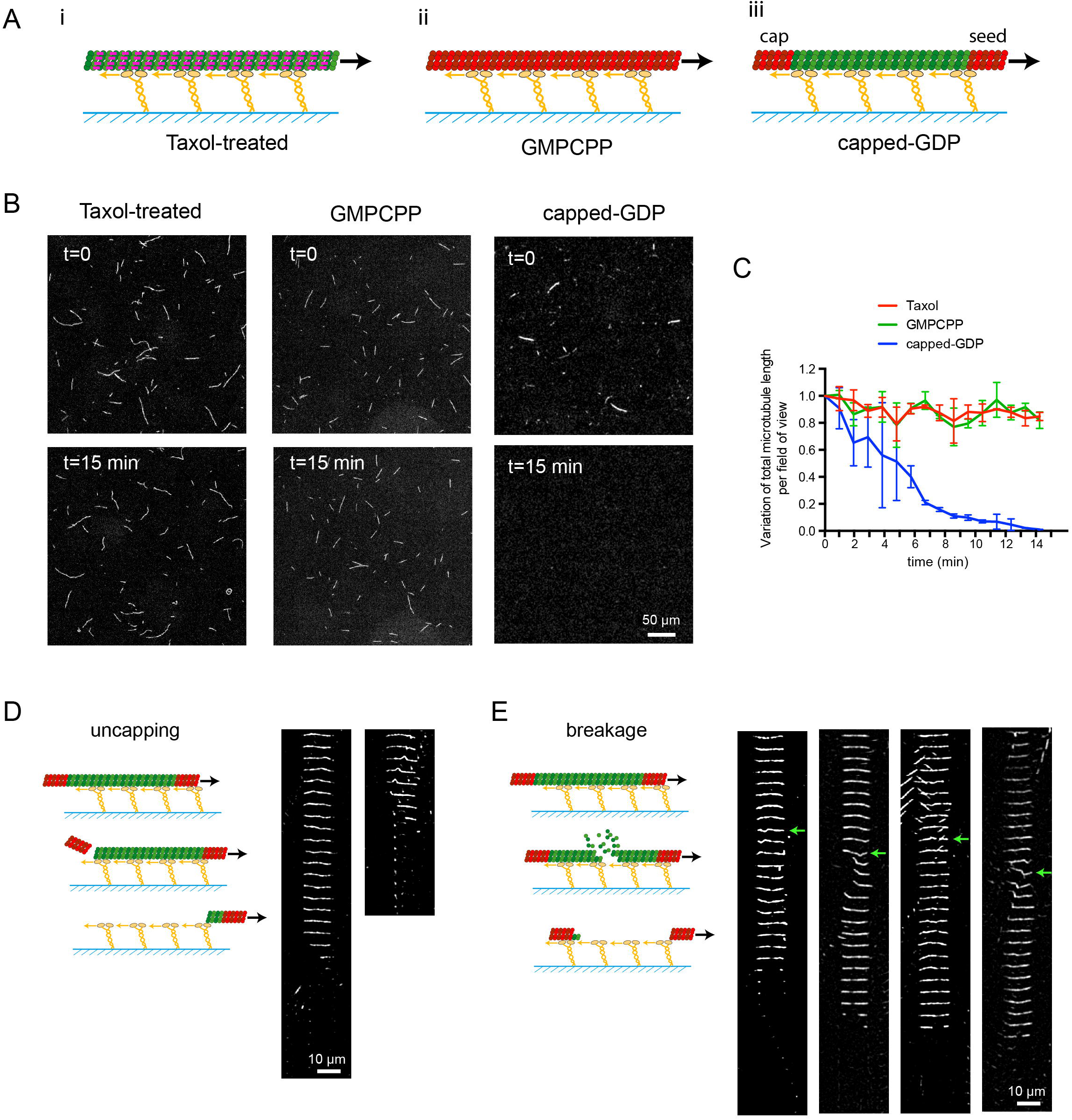
Gliding assays destroy non-stabilized microtubules. A - Schematic representation of the different types of microtubules used in gliding assays. Taxol-treated (purple lines) (i) and GMPCPP-microtubules (red dimers) (ii) are stabilized along their length. Capped-GDP microtubules (iii) are stabilized at their ends (red dimers) but the central part (green dimers) is not. Molecular motors (yellow) are attached to the cover glass (blue), and propel microtubules along the glass surface in the presence of ATP. B - Images of gliding microtubules when motors are activated by the addition of ATP (t=0, top row) and 15 minutes later (t=15 min, bottom row) in the three conditions described in A. Scale bar: 50 μm. C - Quantification of microtubule length variations in the experiments shown in B. The microtubule lengths were measured for all microtubules in the 600-μm-wide fields every minute during 15 minutes. Values were normalized with respect to the initial length (Number of independent experiments N=2, Taxol-treated microtubules n=55; GMPCPP-microtubules n=42; capped-GDP microtubules n=42). D - Schematic representation of microtubule disassembly induced by the loss of the cap. Two examples of microtubules that disassembled from an end during the gliding assay are shown. Images are vertical montages from a movie where images were acquired every 5 s. Scale bar: 10 μm. E - Schematic representation of microtubule disassembly induced by breakage of the central region of the microtubule. Four examples of microtubules that breakage and dissembly during the gliding assay are shown. Images are vertical montages from a movie where images acquired every 5 sec. Green arrows point to bending events that precede the weakening and eventual breakage of the microtubule. Scale bar: 10 μm.

To investigate the mechanism leading to the disassembly of GDP-microtubules, we tracked individual microtubules gliding on kinesin-1, which moves towards the plus ends of microtubules. The majority of the microtubules appeared to depolymerize from their plus end (Figure 1D), consistent with a previous report suggesting that kinesin-1 can trigger the loss of tubulin dimers at the end of stabilized microtubules^23^. In addition, 20% of microtubules broke within the GDP lattice (Figure 1E, movie S3). In some cases breakage events were observed when microtubules encountered other microtubules or when microtubule buckling was observed (Figure 1E, arrows), suggesting that in our assays multiple mechanisms that induce breakage could be in play.

To distinguish between breaking mechanisms we next performed "motility assays” where microtubule-based motility was studied in the opposite geometry, with capped GDP-microtubules attached to the coverslip surface and motors walking along them^24^. Typically, in these assays the microtubules are attached to the surface along their entire length. To attempt to limit these surface interactions (which could interfere with microtubule damage), we used micropatterning to attach microtubule seeds to the surface, leaving the rest of the capped–GDP microtubule unattached to the non-adhesive surface coating^25^ (Figure 2A). We then tested the impact of Klp2, a *Schizosaccharomyces pombe* member of the minus-end-directed kinesin-14 family^26^, on the stability of capped-GDP microtubules. Single Klp2 motors fluorescently labelled with ATTO488 moved processively towards the microtubule minus-end at 120 nm/s, similar to previous reports (Figure 2B)^27^. To limit photo-induced damage we next used non-labelled Klp2 and measured the microtubule lifetime; we observed microtubules breaking and disappearing over the course of an hour (Figure 2C, movie S4). In the absence of kinesins about 88% of microtubules disappeared spontaneously with a lifetime of about 12 min. In the presence of 1 nM Klp2, 96% of microtubules disappeared with a lifetime of about 5 min. Note that the mean lengthes of microtubules for both conditions were not significantly different (16±13 μm and 13±9 μm for the control and at 1 nM Klp2, respectively). While a small percentage of capped-microtubules spontaneously break^11^, we found that Klp2 significantly increased the percentage of broken microtubules (Figure 2D, E). The ability of Klp2 to induce microtubule breakage increased with increasing motor concentration (Figure 2 E, F). At 10 nM Klp2 we measured an average of one breaking event per 14 microns of microtubule (Figure 2F), corresponding in some cases to multiple breaks per microtubule (movie S4). Together our data thus far shows that kinesin motors damage microtubules as they move along them in a manner that leads to microtubule breakage.

**Figure 2.**
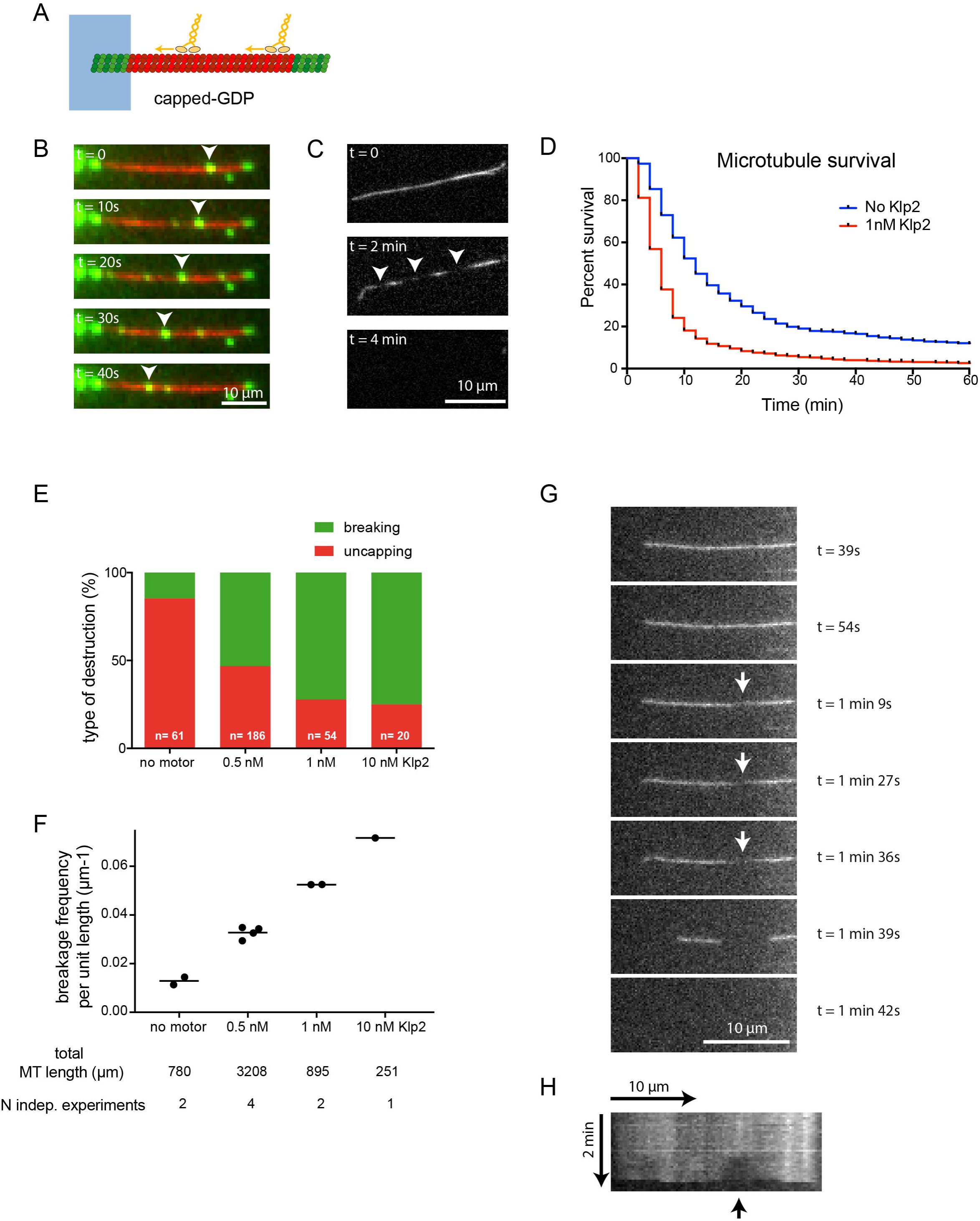
Motility assays destroy non-stabilized microtubules. A - Schematic representation of the motility assay. The microtubule seed (green) is attached to the micropattern (blue), while the shaft (red) and the cap (green) are free. Molecular motors (yellow) walk along the microtubules in the presence of ATP. B - Image sequence showing Klp2 labelled with ATTO488 (green) walking toward microtubule minus-ends. Images were taken every 10 sec. The arrow heads point at a cluster of moving motors. Note that the seed and the cap are also visible in the green channel. Scale bar: 10 μm. C - Image sequence showing the formation of breaks (arrow heads) in the fluorescently-labelled microtubule shaft prior to microtubule disassembly. Motors were not labelled, and images were taken every 2 min. Scale bar: 10 μm. D - Survival curve of microtubules in the absence (blue) or presence (red) of molecular motors. We measured the duration between the addition of molecular motors and the disappearance of microtubules (Number of independent experiments N=2, control n=588, Klp2 n=1130). Data were acquired in the conditions described in C. Pairwaise testing (non-parametric Wilcoxon Mann-Whitney (Wilcoxon rank sum test), two-tailed) of the measured lifetimes gives p-values < 0:0001. E – Bar graph showing the relative proportion of microtubule disassembly after the loss of their cap (red) or a breakage in their central part (green) depending on the concentration of Klp2. Data were acquired in the conditions described in B. F - Graph showing the spatial frequency of microtubule breakage as a function of the concentration of Klp2. Data were acquired in the conditions described in B. G - Image sequence showing the local losses of tubulin dimers in the lattice (arrows) prior to lattice breakage. Scale bar: 10 μm. H - Kymograph showing the evolution over time of fluorescent tubulin intensity along the microtubule shown in G. The arrow points to the progressive enlargment of the section loosing tubulin dimers and the eventual beakage.

In both motility and gliding assays, most breaking events were preceded by a local reduction of microtubule fluorescence, which we hypothesized was due to the loss of tubulin dimers (Figure 1E, Figure 2H arrow). This observations raised the possibility that free tubulin dimers in solution could compensate for this loss by incorporating into the damaged region and protect the microtubule from breakage^28,29^. To test this hypothesis, gliding assays were performed in the presence of free tubulin dimers (Figure 3A). Free tubulin dimers were unlabelled to allow microtubules to be monitored without background fluorescence. Strikingly, capped-microtubules glided on kinesin-1 in the presence of tubulin dimers without any visible destruction (Figure 3B, C, movie S5). Microtubule gliding speed were ~ 500 nm/s in the presence or absence of free dimers (500±75 nm/s and 480±70 nm/s, respectively), showing that free dimers do not interfere with the motor-driven gliding speed of microtubules. The same observation was made in the presence of dynein. Cytoplasmic dynein-1 purified from *Saccharomyces cerevisiae*, like all other dyneins, moves towards the microtubule minus end^30^. Gliding on dyneins in the absence of free dimers led to rapid microtubule disassembly, which could be prevented by the addition of free tubulin dimers (Figure 3D, E, F, movie S6). Similarly, the addition of free dimers in motility assays (Figure 3G) did not interfere with motor movement, but (Figure 3H) fully protected microtubules from motor-induced breakage (Figure 3I, J, movie S7).

**Figure 3.**
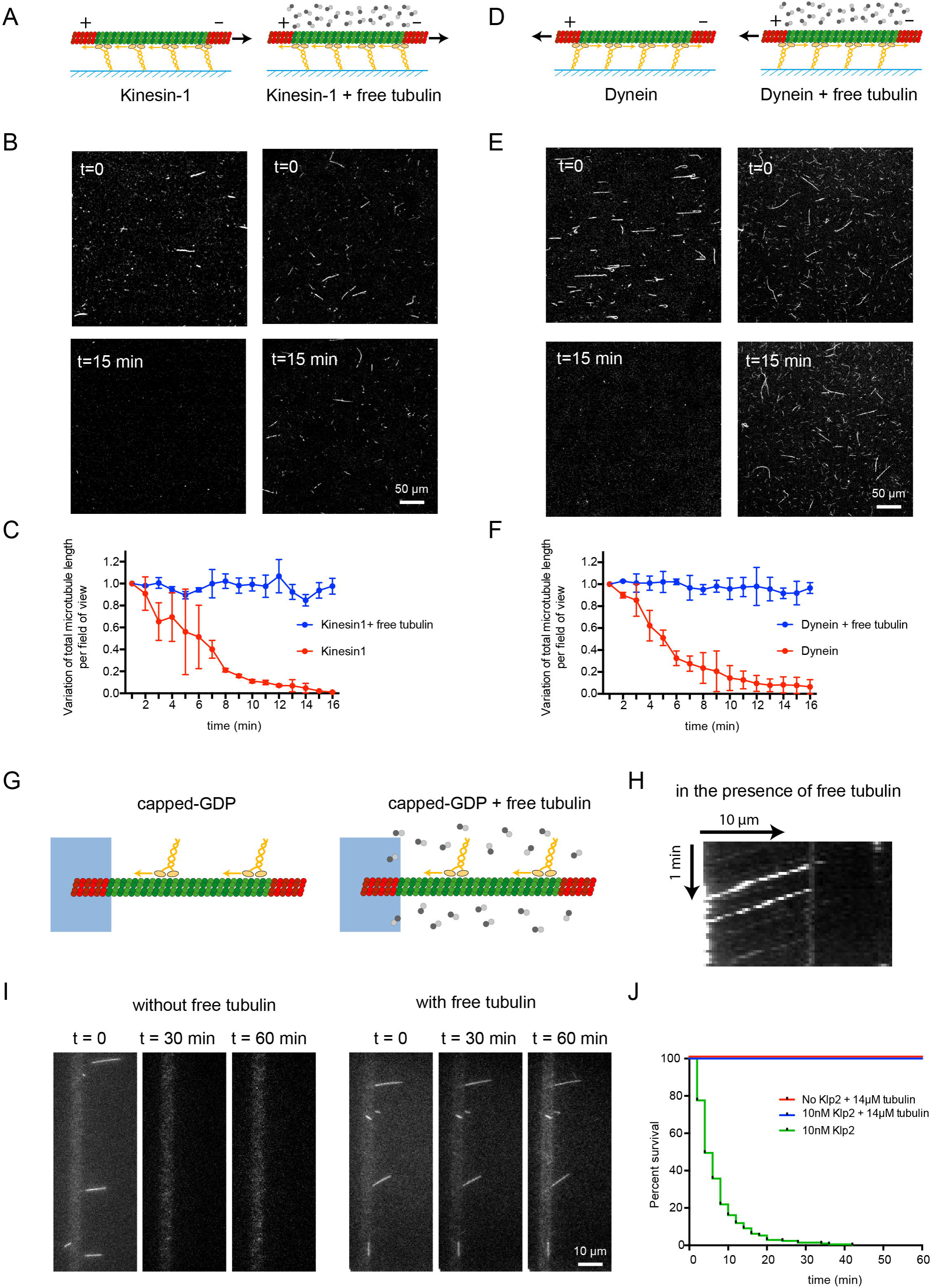
Free tubulin dimers prevent microtubule destruction by kinesin and dynein. A - Schematic representation of the gliding assay of capped-GDP microtubules on kinesin-1 in the absence (left) or presence (right) of free non-labelled tubulin dimers. B - Images of gliding microtubules when motors are activated by the addition of ATP (t=0, top row) and 15 minutes later (t=15 min, bottom row) in the two conditions described in A. C - Quantification of microtubule length variations in the experiments shown in B. The microtubule lengths were measured for all microtubules in the 600-μm-wide fields every minute during 15 minutes. Values were normalized with respect to the initial length. Data in the absence of free tubulin dimers are shown in red and correspond to the data shown in figure 1C, those in the presence of free dimers are shown in blue. Values were normalized with respect to the initial intensity (Number of independent experiments N=2, without free tubulin n=42, with free tubulin n=74). D,E,F – Same as A, B and C in the presence of yeast dynein instead of kinesin-1. (N=2, without free tubulin n=24, with free tubulin n=12). G - Schematic representation of the motility assay in the absence (left) and presence (right) of free non-labelled tubulin dimers. H - Kymograph showing the position of fluorescent-Klp2 position along a microtubule over time. The presence of free dimer did not interfere with the ability of the motor to walk towards microtubule minus ends. I - Image sequences show microtubules after the addition of Klp2 motors in the absence (left) or presence (right) of free tubulin dimers. Images were taken every 2 min. Scale bar: 10 μm. J - Survival curve of microtubules in the presence of molecular motors (green), in the presence of molecular motors and free tubulin dimers (blue) or in the absence of motors and presence of free tubulin dimers (red). Data where acquired in the conditions described in I. (N=1; No motor, n=108; 10nM Klp2 with free tubulin, n=70; 10nM Klp2 without free tubulin, n=212).

To directly visualize microtubule repair in response to motor-induced destruction, we performed a two-color assay in which polymerized tubulin and free tubulin dimers were labelled with distinct fluorophores. This allowed us to visualize tubulin incorporation at damaged sites^29^ (Figure 4A). In a kinesin-1 gliding assay in the presence of free green-tubulin dimers, red-fluorescent capped-GDP microtubules displayed micrometer-long green stretches along their shaft 30 minutes after the initiation of the gliding assay (Figure 4B). These two colored microtubules could also be seen moving along the kinesin-1 surface (movie S8). Similar two-colored microtubules were observed in dynein microtubule gliding assays performed in the same manner (Figure 4C). By varying the duration of the gliding step during the 30-minute assay (step 2 in Figure 4A) we found that the size of the repaired sites remained constant, but that their frequency along the shaft increased regularly (Figure 4D). This suggests that motors continuously generate new sites of damage. Note that the presence of repaired sites in microtubules which did not glide because of the absence of ATP during the 30 minutes of the assay (“time 0” in figure 4D) was consistent with our recent observation that microtubule lattice has an intrinsic turnover rate^11^. In gliding microtubules, the increased frequency of repair sites corresponded to the addition of the intrinsic and the motor-induced contributions. The spatial frequency of the repaired sites was lower in the dynein gliding assay compared to the kinesin assay, perhaps due to differences in the force production and velocity of these two motors. Yeast dynein moves about 10-fold slower than kinesin-1 and also has a slightly lower stall force^31^.

**Figure 4.**
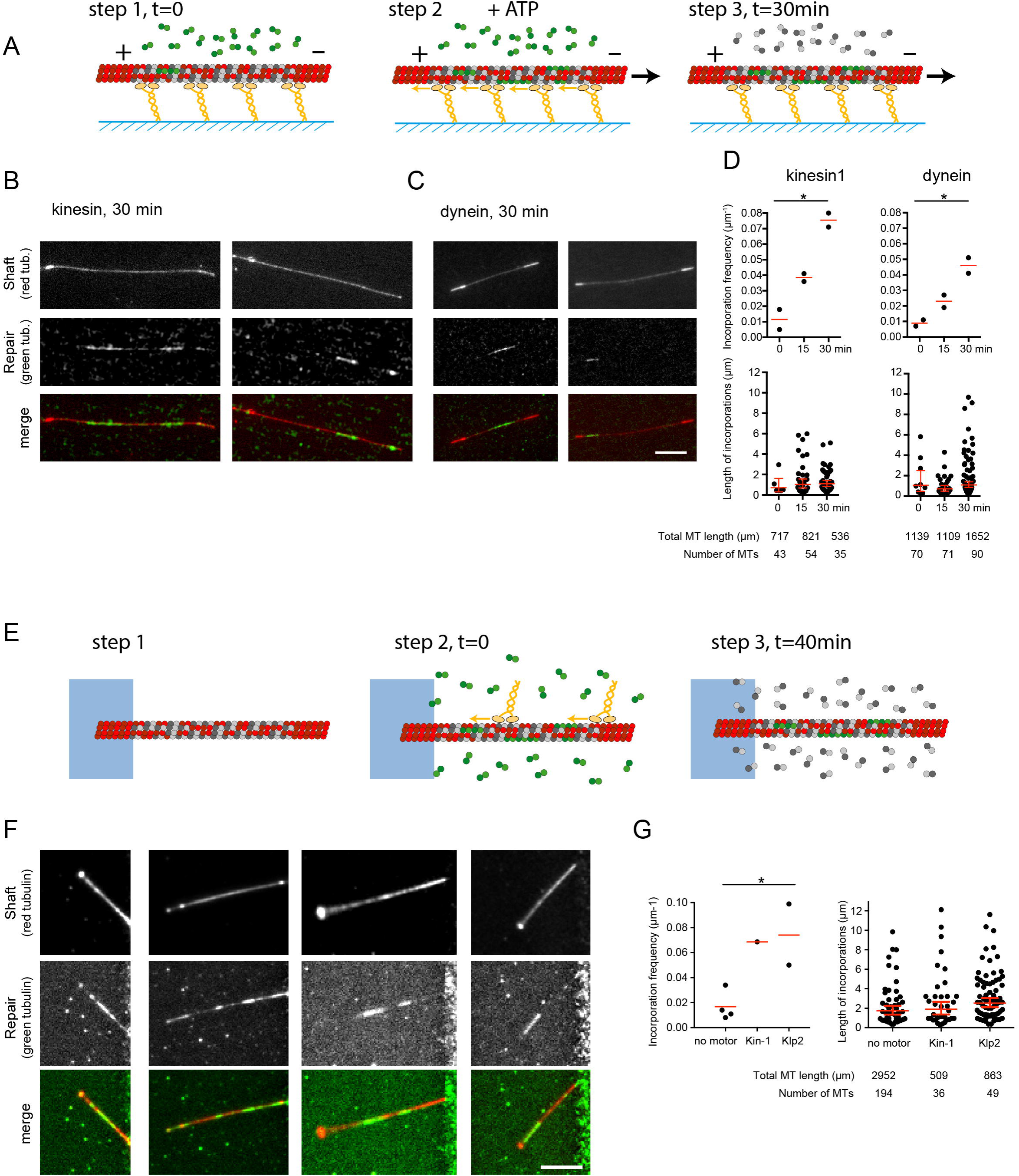
Molecular motors catalyze tubulin exchange in the microtubule lattice. A - Schematic representation of the three sequences during gliding assays of capped-GDP microtubules in the presence of free fluorescent tubulin dimers. All experiments lasted 30 minutes. Only the duration of the gliding step was varied. Step1 corresponds to the loading of capped-GDP microtubules and free green tubulin dimers on the layer of motors. Step 2 corresponds to the initiation of gliding by the addition of ATP. In the 0-minute condition, ie without gliding, step 2 was skipped and ATP was not added. In the 15-minute condition, step 2 was initiated 15 minutes after step 1. In the 30-minute condition, step 2 immediately followed step 1. In all cases, fluorescent tubulin dimers were removed after 30 minutes to measure the incorporation of free tubulin dimers. B, C – Images show the microtubule shaft (low intensity) with associated cap and seed (high intensity) (top row), the incorporation of free tubulin dimers (middle row), and the overlay of the two signals (bottom row) at the end of the gliding assay (step 3). In these examples, microtubules glided on kinesin-1 (B) or dynein (C) for 30 minutes. Scale bar: 10 μm. D – Quantifications of the spatial frequency (top) and size (bottom) of the incorporation sites upon gliding on kinesin-1 (left) or dynein (right) depending on the duration of the gliding step. Data were acquired during two independent experiments. Each experiment allowed the measurement of a single value of incorporation frequency over several microtubules. Red bars show geometrical means with 95% confidence interval. The table indicates the number and total length of microtubules that were measured in each condition. E - Schematic representation of the three sequences during motility assays on capped-GDP microtubules in the presence of free fluorescent tubulin dimers. Step 1 is the growth and capping of microtubules. Step 2 corresponds to the addition of molecular motors and free fluorescent tubulin dimers. Step 3 corresponds to the removal of motors and replacement of fluorescent tubulin dimers by non-labelled tubulin dimers 40 minutes later. F - Images show the microtubule shaft (top row), the incorporation of free dimers (middle row), and the overlay of the two signals (bottom row) at the end of the motility assay. G - Quantification of the spatial frequency (top) and size (bottom) of the incorporation sites following the motility of kinesin-1 (10 nM) or Klp2 (10 nM). Data were acquired during four independent experiments for the control conditions without motors, a single experiment with kinesin-1 and two independent experiments with Klp2. Each experiment allowed the measurement of a single value of incorporation frequency over several microtubules. Red bars show geometrical means with 95% confidence interval. The table indicates the number and total length of microtubules that were measured in each condition.

The integration of free tubulin dimers into the microtubule shaft could also be visualized in motility assays using a similar experimental strategy (Figure 4E). Clear stretches of green tubulin dimers along the shaft of red microtubules were present on 80% (n=85) of microtubules after kinesin-1 or Klp2 moved processively along them (Figure 4F). The frequency of the repaired sites was similar to those measured in gliding assays, but the size of the repaired sites was larger (Figure 4G). This data as well as the kinetics of microtubule destruction in the absence of free tubulin dimers suggests that freely fluctuating microtubules are more amenable to lattice disruption than those bound to a glass slide. Together these results show that walking motors can break the microtubule lattice, catalysing the incorporation of free tubulin dimers, which heal the lattice and protect it from disassembly. From this data, we infer that components of the microtubules that are supporting motor trafficking are continuously renewed.

Actin-based myosin motors, which contract the actin network through pulling forces, can also destroy their tracks^32,33^. Therefor, our work suggests that the coupling between working motors and cytoskeleton disassembly is a general feature that allows remodelling of the cytoskeletal filament that is under strain. Importantly, under biochemical conditions that allow the self-repair of the constrained microtubule lattice, the incorporation of new GTP-tubulin dimers generates rescue sites along microtubules, increasing their lifespan^34–36^. This raises the possibility that dynein and kinesin motors may trigger the selective repair and stabilization of the microtubules they are walking on.

## Methods

### Microtubule gliding assay

In vitro gliding assays were performed using flow chamber with dimensions of 2 × 10 × 0.07 mm (W × L × H), which were assembled from 10 × 10 mm and 24 × 30 mm cover glasses (Thermo Scientific) with double-sided tape as the spacer. Anti-GFP antibody (Invitrogen) at 0.2 mg mL-1 (5 μL) was applied to the flow chamber for gliding assays with kinesins. The flow chamber was washed with 5 μL of 1% w/v BSA in HKEM (10 mM HEPES pH=7.2, 50 mM KCl, 1 mM EGTA, 5 mM MgCl_2_). Next 5 μL of 200 nM GFP-tagged kinesin-1 or 3 μL of 10 nM yeast dynein in wash buffer (10 mM HEPES, 16 mM PIPES (pH 6.8), 50 mM KCl, 5 mM MgCl_2_, 1 mM EGTA, 20 mM DTT, 3 mg/ml glucose, 20 μg/ml catalase, 100 μg/ml glucose oxidase, and 0.3% BSA) was introduced for 3 min and washed with 10 μL wash buffer. 10 μL of microtubules in HKEM was then introduced and incubated for 1 min, followed by extensive washing with wash buffer. Finally, 10 μL of ATP buffer (10 mM HEPES, 16 mM PIPES (pH 6.8), 50 mM KCl, 5 mM MgCl_2_, 1 mM EGTA, 20 mM DTT, 3 mg/ml glucose, 20 μg/ml catalase, 100 μg/ml glucose oxidase, 0.3% BSA, 2.7 mM ATP, and 0.2% Methyl cellulose) was added into the flow chamber prior to imaging. In the experiment with taxol-stabilized microtubules, washing buffers and ATP buffers contained 10 μM taxol. In the experiments in the presence of free tubulin, 14 μM of unlabelled tubulin was added to the ATP buffer.

### Microtubule motility assay

Micropatterning of glass slides and fabrication of the microfluidic circuitry are described in the Supplementary methods section.

Microtubule seeds were prepared at 10 μM tubulin concentration (20% ATTO651-labeled and 80% biotinylated tubulin) in BRB80 (80mM PIPES pH=6.8, 1mM MgCl_2_, 1mM EGTA) supplemented with 0.5 mM GMP-CPP at 37°C for 1 h. The seeds were incubated with 1 μM Taxotere (Sigma) at room temperature for 30 min and were then sedimented by centrifugation at 30°C for 15min at 100,000 x g and resuspended in BRB80 supplemented with 0.5 mM GMP-CPP and 1 μM Taxotere. Microtubule seeds were stored in liquid nitrogen and quickly warmed to 37°C before use.

The Poly-DiMethyl-Siloxane (PDMS) chip was placed on a micropatterned cover glass and fixed on the microscope stage. The chip was perfused with neutravidin (50 μg/ml in HKEM; Pierce), then washed with HKEM supplemented with 0.2% BSA and passivated for 1 min with PLL-g-PEG (Pll 20K-G35-PEG2K, Jenkam Technology) at 0.1 mg/ml in 10 mM Na-Hepes (pH 7.4), and washed again with HKEM plus 0.2% BSA. Microtubule seeds were flowed into the chamber at high flow rates perpendicularly to the micropatterned lines to ensure proper orientation of the seeds. Non-attached seeds were washed out immediately using HKEM supplemented with 0.2% BSA. Seeds were elongated with a mix containing 20 μM tubulin (20% labeled with ATTO-647) in 10 mM HEPES, 16 mM PIPES (pH 6.8), 50 mM KCl, 5 mM MgCl_2_, 1 mM EGTA, 20 mM DTT, 3 mg/ml glucose, 20 μg/ml catalase, 100 μg/ml glucose oxidase, 0.3% BSA and 0.25% methyl cellulose (1500 cp, Sigma) for 20 min. GMPCPP caps were grown by supplementing the before-mentioned buffer for with 0.5 mM GMPCPP (Jena Bioscience) instead of GTP and using 14 μM tubulin (100% labeled with a ATTO-488 or ATTO-565 fluorophore) for 15 min. To monitor microtubule survival in the absence of free tubulin, the same buffer as for seed elongation was used, but without free tubulin.

### Visualization of free dimer incorporation

To visualize incorporation of free tubulin dimers into the microtubule shaft during gliding, the glass surface was passivated with 1% w/v BSA and 1% w/v pluronic F-127. Unlabelled SNAP-kinesin1-His was used instead of GFP-kinesin1-His. Green fluorescent dimers tend to attach to the layer of motors during the gliding assay, hindering the detection of the few dimers that may incorporate into the lattice. To limit this effect, the last coating step was performed with wash buffer containing 14 μM unlabelled tubulin. Capped-GDP microtubules were assembled in a single red colour, but with a lower proportion of fluorescent dimers (3%) in the central GDP part compared to stabilized ends (20% labelled tubulin). Fluorescence in the central part of the microtubule was required to visualize capped-microtubules and distinguish them from individual seeds. The gliding of these capped-microtubule was initiated with ATP buffer containing 14 μM tubulin (20% labelled with green fluorophore) and recorded for 30 min. In the control experiment, ATP was not included in the reaction mixture. At the end of the gliding experiments, labelled tubulin was washed out and replaced by new ATP buffer with 7 μM unlabelled tubulin in order to visualize green dimer incorporation into the capped-microtubules without triggering microtubule destruction due to dimer washout.

To visualize incorporation of free tubulin dimers into the microtubule shaft during motility assays, capped-microtubules were exposed to 14 μM of tubulin (100% labeled, green or red fluorescent) in 10 mM HEPES, 16 mM PIPES (pH 6.8), 50 mM KCl, 5 mM MgCl_2_, 1 mM EGTA, 20 mM DTT, 3 mg/ml glucose, 20 μg/ml catalase, 100 μg/ml glucose oxidase, 0.3% BSA and 0.25% methyl cellulose (1500 cp, Sigma), in the absence or presence of motors, for 40 min before replacing it with elongation buffer in which fluorescent tubulin dimers were replaced by 10 μM unlabeled tubulin in order to protect microtubules from disassembly without perturbing the imaging process with background fluorescence.

### Imaging

Large scale measurement of microtubule lifetime during gliding assays were performed by illuminating the sample with a SOLA Light engine (Lumencor) and visualised with an epifluorescence microscope (Eclipse Ti2, Nikon) using a S Plan Fluor 20X objective lens (Nikon). The microscope stage was kept at 37°C using a warm stage controller (LCI). Images and movies were captured using a CMOS camera (Hamamatsu) every 5 sec for 15 min.

Individual microtubule destruction or repair events during gliding and motility assays were visualized using an objective-based TIRF composed of a Nikon Eclipse Ti, an azimuthal ilas2 TIRF illuminator (Roper Scientific), a 60x NA 1.49 TIRF objective lens followed by a 1.5X magnification lens and an Evolve 512 camera (Photometrics). The microscope stage was kept at 37°C using a heated stage controller (LCI). Excitation was achieved using 491 and 565 nm or 642 nm lasers (Optical Insights). Time-lapse recording was performed using Metamorph software (version 7.7.5, Universal Imaging). Images were taken every second.

Large scale measurement of microtubule lifetime during motility assays were performed by acquiring images every 2 min for 1 hour. To detect microtubule breakage, images were taken every 3 sec until microtubule disappeared. To visualize free tubulin dimer incorporation into polymerized microtubules, images were taken every sec for 15 sec at the end of the assay so that images could be averaged to improve the signal to noise ratio.

### Image processing and measurements

For gliding assays, microtubule lifetime was quantified by measuring the variation of the microtubule length over time by using the line tool of Adobe Photoshop CC2018. The quantification of the free tubulin incorporation along the microtubule shaft was done using a line scan in Image J after the following image treatment. We arbitrary selected three frames at different time points for the green and red channels (incorporated tubulin is green and the microtubule is red). Overlaid images were generated using an Adobe Photoshop original macro for each channel by cancelling the displacement of the microtubule during gliding. To generate the image overlay, the Photoshop layer mode “darken mode” was applied to each image in order to reduce background noise. Finally, the background of the overlaid image was subtracted by Image J (Rolling ball radius 10.0 pixels). The incorporation was defined as signal 1.5 times higher than background (see Supplementary Fig. S1).

For motility assays, movies were processed to improve the signal/noise ratio. Background was substracted with a dedicated plug-in from imageJ, version 1.47n5). Microtubule lifetime was quantified by manually counting microtubule number at each frame, every 2 min for 1 hour. To quantify the incorporation of free tubulin dimers, line scans were used to measure fluorescence intensity along the microtubule. Incorporation was counted when a spot perfectly aligned with the microtubule lattice and the fluorescence intensity showed a peak at least 2-fold higher than the background fluorescence.

## References

1. VanBuren, V., Cassimeris, L. & Odde, D. J. Mechanochemical model of microtubule structure and self-assembly kinetics. Biophys. J. 89, 2911–2926 (2005).

2. Vale, R. D., Reese, T. S. & Sheetz, M. P. Identification of a novel force-generating protein, kinesin, involved in microtubule-based motility. Cell 42, 39–50 (1985).

3. Paschal, B. M. & Vallee, R. B. Retrograde transport by the microtubule-associated protein MAP 1C. Nature 330, 181–183 (1987).

4. Lye, R. J., Porter, M. E., Scholey, J. M. & McIntosh, J. R. Identification of a microtubule-based cytoplasmic motor in the nematode C. elegans. Cell 51, 309–318 (1987).

5. Schnitzer, M. J. & Block, S. M. Kinesin hydrolyses one ATP per 8-nm step. Nature 388, 386–390 (1997).

6. Drummond, D. R. Regulation of microtubule dynamics by kinesins. Semin. Cell Dev. Biol. 22, 927–934 (2011).

7. Hibbel, A. et al. Kinesin Kip2 enhances microtubule growth in vitro through length-dependent feedback on polymerization and catastrophe. Elife 4, 1–11 (2015).

8. Laan, L. et al. Cortical Dynein Controls Microtubule Dynamics to Generate Pulling Forces that Position Microtubule Asters. Cell 148, 502–514 (2012).

9. Hunter, A. W. et al. The kinesin-related protein MCAK is a microtubule depolymerase that forms an ATP-hydrolyzing complex at microtubule ends. Mol. Cell 11, 445–57 (2003).

10. Arellano-Santoyo, H. et al. A Tubulin Binding Switch Underlies Kip3/Kinesin-8 Depolymerase Activity. Dev. Cell 42, 37–51.e8 (2017).

11. Schaedel, L. et al. Lattice defects induce microtubule self-renewal. *bioRxiv* (2017). doi: 10.1101/249144

12. Peet, D. R., Burroughs, N. J. & Cross, R. A. Kinesin expands and stabilizes the GDP-microtubule lattice. Nat. Nanotechnol. (2018). doi:10.1038/s41565-018-0084-4

13. Shima, T., Morikawa, M., Kaneshiro, J. & Kambara, T. Kinesin-binding triggered conformation switching of microtubules contributes to polarized transport. 1–20 (2018).

14. Hancock, W. O. The Kinesin-1 Chemomechanical Cycle: Stepping Toward a Consensus. Biophys. J. 110, 1216–1225 (2016).

15. Cianfrocco, M. A., DeSantis, M. E., Leschziner, A. E. & Reck-Peterson, S. L. Mechanism and Regulation of Cytoplasmic Dynein. Annu. Rev. Cell Dev. Biol. 31, 83–108 (2015).

16. Visscher, K., Schnltzer, M. J. & Block, S. M. Single kinesin molecules studied with a molecular force clamp. Nature 400, 184–189 (1999).

17. Carter, N. J. & Cross, R. A. Mechanics of the kinesin step. Nature 435, 308–312 (2005).

18. Mickolajczyk, K. J. et al. Kinetics of nucleotide-dependent structural transitions in the kinesin-1 hydrolysis cycle. Proc. Natl. Acad. Sci. 112, E7186–E7193 (2015).

19. Kawaguchi, K. Energetics of kinesin-1 stepping mechanism. FEBS Lett. 582, 3719–3722 (2008).

20. Block, S. M. Kinesin motor mechanics: Binding, stepping, tracking, gating, and limping. Biophys. J. 92, 2986–2995 (2007).

21. VanBuren, V., Odde, D. J. & Cassimeris, L. Estimates of lateral and longitudinal bond energies within the microtubule lattice. Proc. Natl. Acad. Sci. 99, 6035–6040 (2002).

22. Howard, J., Hudspeth, A. J. & Vale, R. D. Movement of microtubules by single kinesin molecules. Nature 342, 154–158 (1989).

23. Dumont, E. L. P., Do, C. & Hess, H. Molecular wear of microtubules propelled by surface-adhered kinesins. Nat. Nanotechnol. 10, 1–4 (2015).

24. Vale, R. D. et al. Direct observation of single kinesin molecules moving along microtubules. Nature 380, 451–453 (1996).

25. Portran, D., Gaillard, J., Vantard, M. & Théry, M. Quantification of MAP and molecular motor activities on geometrically controlled microtubule networks. Cytoskeleton (Hoboken). 70, 12–23 (2013).

26. Carazo-Salas, R. E., Antony, C. & Nurse, P. The kinesin Klp2 mediates polarization of interphase microtubules in fission yeast. Science 309, 297–300 (2005).

27. Braun, M., Drummond, D. R., Cross, R. A. & McAinsh, A. D. The kinesin-14 Klp2 organizes microtubules into parallel bundles by an ATP-dependent sorting mechanism. Nat. Cell Biol. 11, 724–30 (2009).

28. Dye, R. B., Flicker, P. F., Lien, D. Y. & Williams, R. C. End-stabilized microtubules observed in vitro: stability, subunit, interchange, and breakage. Cell Motil. Cytoskeleton 21, 171–86 (1992).

29. Schaedel, L. et al. Microtubules self-repair in response to mechanical stress. Nat. Mater. 14, 1156–1163 (2015).

30. Reck-Peterson, S. L. et al. Single-Molecule Analysis of Dynein Processivity and Stepping Behavior. Cell 126, 335–348 (2006).

31. Gennerich, A., Carter, A. P., Reck-Peterson, S. L. & Vale, R. D. Force-Induced Bidirectional Stepping of Cytoplasmic Dynein. Cell 131, 952–965 (2007).

32. Haviv, L., Gillo, D., Backouche, F. & Bernheim-Groswasser, A. A cytoskeletal demolition worker: myosin II acts as an actin depolymerization agent. J. Mol. Biol. 375, 325–30 (2008).

33. Reymann, A.-C. et al. Actin network architecture can determine myosin motor activity. Science (80-.). 336, (2012).

34. Aumeier, C. et al. Self-repair promotes microtubule rescue. Nat. Cell Biol. 18, (2016).

35. de Forges, H. et al. Localized Mechanical Stress Promotes Microtubule Rescue. Curr. Biol. 26, 3399–3406 (2016).

36. Vemu, A. et al. Severing enzymes amplify microtubule arrays through lattice GTP-tubulin incorporation. Science (80-.). 361, eaau1504 (2018).

